# Voltage Imaging with Periodic Structured Illumination

**DOI:** 10.1101/2025.07.24.666645

**Authors:** Forest Speed, Alec Teel, Gregory L. Futia, Diego Restrepo, Emily A. Gibson

**Affiliations:** Department of Bioengineering, University of Colorado Anschutz Medical Campus, Aurora, CO 80045, USA; Department of Cell and Developmental Biology, University of Colorado Anschutz Medical Campus, Aurora, CO 80045, USA

## Abstract

We utilize periodic structured illumination with pseudo-HiLo (pHiLo) image reconstruction for in vivo voltage imaging. We demonstrate reduced signal from out-of-focus cells, that contaminates voltage activity for in-focus cells of interest, with pHiLo compared to traditional widefield recordings taken with uniform illumination and pseudo-widefield (pWF) reconstructions. We discuss tradeoffs between signal-to-background ratio, signal-to-noise ratio and temporal resolution for pHiLo in the context of high-speed voltage imaging in awake mice.

## 1. Introduction

Fluorescence microscopy with genetically encoded calcium indicators (GECIs) and genetically encoded voltage indicators (GEVIs) provides a toolset for studying neuronal activity in behaving animals [1-6]. Cells that express GECIs exhibit fluorescent transients in response to changes in cytoplasmic calcium concentration that occur after an action potential. In contrast, cells expressing GEVIs exhibit fluorescence changes that directly follow cellular membrane potential. Because GEVIs are membrane localized, the total fluorescence signal is inherently reduced compared to the signal from GECIs which fill the cytosol. Furthermore, the sub-millisecond response kinetics of GEVIs necessitate much higher frame rates for action potential detection, decreasing available exposure time and increasing signal variance due to Poisson noise. Due to the high frame rate requirement of voltage imaging, calcium imaging is more commonly used for in vivo functional imaging applications. However, intracellular calcium concentration is not analogous to membrane potential. Slow calcium efflux kinetics necessitate computational inference of high-frequency spike trains [1]. This advantage of GEVIs necessitates their use in accurately measuring neural activity.

The main optical approaches for neural recording are 2-photon fluorescence microscopy (2P, Figure 1A) and widefield fluorescence microscopy (WF, Figure 1B) [7-10]. 2P microscopy is more commonly used for in vivo calcium recording because of the inherent ability to provide 3D optical sectioning and penetrate deeper into tissue. With 2P approaches, the data that is recorded only consists of fluorescence emitted from cells in the focal plane of the microscope as well as accompanying noise from the readout electronics (read noise) and photon noise. For N photons collected from in-focus neurons, the standard deviation of the 2P recorded signal caused by photon noise is approximated as √N. 2P microscopes rely on mechanical scanning components that places an upper bound on achievable image frame rates for 2P, typically up to 80 Hz.

**Fig. 1.**
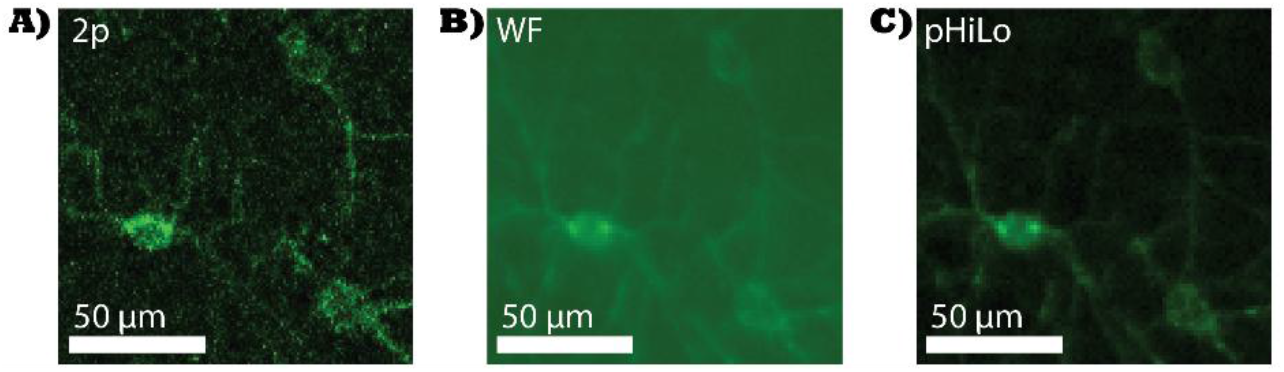
Comparison of 2-photon (2P), Widefield (WF) and Pseudo-HiLo (pHiLo) images of NDNF interneurons expressing Voltron2_552_ in the visual cortex of an awake animal. **A**. 2P image taken with 2 μs dwell time, 12 Hz frame rate, and 3-frame averaging. **B**. WF image taken with a 2 ms exposure time. **C**. pHiLo image taken with a 2 ms exposure time. The structured illumination reconstruction of pHiLo combines the benefits of contrast enhancement of 2P and high frame rate capabilities of WF.

Strategies to adapt 2p systems for use with voltage indicators require complex and expensive optoelectronics [7, 9, 11-14]. High-speed 2P microscopy has been demonstrated by employing multiple focal spots for parallel acquisition, however this is ultimately limited by photodamage thresholds. Alternatively, fast acousto-optic deflectors have been employed to rapidly scan between different regions of interest, but the number of cells that can be recorded in a serial manner are limited [15]. To reach comparable voltage imaging capabilities as widefield imaging systems, significant improvements must be made to the brightness and sensitivity of 2p optimized voltage indicators [13, 15]. Furthermore, the optoelectronic complexity of 2p voltage imaging systems restricts their use to benchtop systems that cannot easily be implemented with miniature microscope technology. The primary motivation for our utilization of widefield imaging and periodic structured illumination is develop techniques compatible with existing miniscope devices developed for freely behaving animals [16, 17].

Widefield techniques enable considerably high frame rates (up to 2 kHz) and larger field of view (FOV) than 2p but lack inherent optical sectioning and deep tissue penetration capability. In a typical widefield recording, the entire sample is excited using uniform illumination and the fluorescence emitted from both in-focus and out-of-focus sources is collected by an image sensor. The recorded information can be decomposed into three main components: fluorescence emitted from in-focus cells, fluorescence emitted from out-of-focus cells, and the combination of read and shot noise [18, 19]. For most applications, signal variance due to shot noise is multiple orders of magnitude larger than from read noise. For B photons collected from out-of-focus features, the widefield shot noise approximation is √(N+B) as the total photons detected include those from both in-focus signal and out-of-focus background [15]. While the rate of widefield microscopy is not limited by the need for mechanical scanning, it is limited by reduced contrast between the in-focus signal and out-of-focus background (Figure 1B). The time dependent fluorescence signal from out-of-focus cells contributes to the background temporal fluctuations that can contaminate intensity time courses extracted from in-focus cells [18, 20-25]. It is therefore beneficial for widefield systems to have a strategy for background identification and removal. Periodic structured illumination with image reconstruction offers one such strategy. In contrast to traditional widefield recordings that use uniform illumination, structured illumination uses a regularly spaced grating pattern [26-30]. The modulation contrast induced by the grating pattern is highest for in-focus information, enabling its separation from the out-of-focus background. In traditional OS-SIM, the reconstruction relies on acquiring three separate images with the illumination shifted by 0, 120, and 240 degrees: I_0_, I_120_, I_240_. The in focus (i.e. modulated) signal is isolated using square law detection (Eq. 1) [27].

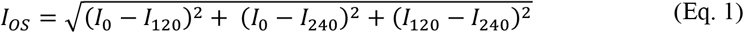

However, the utilization of traditional structured illumination reconstruction techniques is fundamentally limited in the context of time-dependent voltage imaging as they rely on squared-subtractions between consecutive frames to evaluate grating contrast [31-33]. One alternative method is HiLo imaging [21, 28, 34]. In HiLo, images are collected with alternating structured illumination (I_SI_) and uniform illumination (I_UI_). Modulation contrast imparted by the grating pattern is extracted from a difference image (I_DI_). Before computing I_DI_, the frames are each normalized by dividing the intensity at each pixel by a local average obtained via Gaussian low-pass filtering at a cutoff lower than the grating frequency [34]. The effect of the normalization stage is that global intensity differences do not cause reconstruction artifacts. I_DI_ is then obtained by subtracting the normalized uniform illumination image from the normalized structured illumination image [34]. In the ideal (noise-free) case, pixels in I_DI_ that correspond to locations composed of only out-of-focus fluorescence will be zero. A spatially varying modulation contrast map is then obtained through analysis of local contrast in the difference image [34]. In-focus low spatial frequency information (I_Lo_) is obtained by applying a Gaussian low-pass filter (using the grating frequency as cutoff) to the product of the modulation contrast map and I_UI_ [34]. High-frequency information (I_Hi_) is obtained by applying a high-pass filter to I_UI_ with a cutoff at the grating frequency [34]. The HiLo reconstruction is then obtained by combining I_Hi_ and I_Lo_ after scaling I_Lo_ by a user-define parameter, α, for continuity of spatial frequency (I_HiLo_ = I_Hi_ + αI_Lo_) [34].

In previous work, we developed a HiLo-based reconstruction technique for data recorded using only periodic structured illumination (pHiLo) [26]. Our motivation for pHiLo stems from the development of microLED stripe arrays designed to provide structured illumination excitation for miniature microscope applications with freely behaving mice [16, 17]. To obtain a pHiLo reconstruction, data is first recorded using periodic structured illumination with the grating pattern shifting by 120 degrees between consecutive frames. A 3-frame boxcar average is then applied to each pixel of the image stack to generate a pseudo-widefield (pWF) stack. Traditional HiLo reconstruction is then performed with the raw structured illumination stack and pWF stacks serving as inputs for I_SI_ and I_UI_, respectively [26]. Recording GCaMP8f activity in the hippocampus with periodic structured illumination, we found that pHiLo reconstruction provided significant benefits compared to OS-SIM. While both reconstruction techniques adequately removed contamination from out-of-focus background components, pHiLo exhibited improved signal-to-noise ratio (SNR) and reduced distortion due to motion encountered with in vivo imaging compared to OS-SIM [26].

In this work, we implement the pHiLo technique for recording voltage activity with periodic structured illumination in head-fixed mice expressing Voltron2 [2]. While other techniques have been demonstrated for widefield voltage imaging with patterned illumination [20, 35, 36], pHiLo uses only periodic structured illumination images that can be generated with microLED striped emitters [17, 37]. By utilizing pHiLo for voltage imaging in vivo, we demonstrate a simple and reproducible methodology to obtain optically sectioned reconstructions of voltage dynamics at widefield frame rates using only periodic structured illumination.

## 2 Experimental Methods

### 2.1 Imaging System

A system diagram for the one photon upright microscope with patterned illumination is shown in Figure 1. A digital micromirror device (Texas Instruments, DLP6500/DLPC900) patterns light emitted from a 5W 532nm laser (Coherent, Verdi V5). The beam is focused onto the sample with 20-100 mW/mm^2^ of intensity using a 16x/0.8 NA long working distance (LWD) objective (Nikon, CFI75) and emitted fluorescence is separated using a shortpass 567 nm dichroic mirror (Thorlabs, DMSP567L). Separated fluorescence passes through a long pass emission filter (Semrock, BLP01-568R-25) and 0.35x magnification coupling (Olympus, U-TV0.35XC) before it is recorded by a back-illuminated sCMOS (Hamamatsu Orca-Fusion BT, C15440-20UP) with ∼95% quantum efficiency at 581 nm (peak emission wavelength of Voltron2_552_). The camera operates in SYNCREADOUT mode with external triggers provided by the DMD. The grating patterns used in this work ranged from 14 μm to 25 μm. The mean fluorescence collected from regions of information (ROIs) around in-focus cells evaluated in this work was at least 500 photons per frame per pixel under uniform illumination (Figure 5).

Two-photon images were acquired with an upright two photon microscope (3i, VIVO Multiphoton RS+ Movable Objective Microscope Workstation) with a tunable wavelength laser (SpectraPhysics, MaiTaiHP) at 1030 nm and 25x/1.05NA objective (Olympus, XLPlanN).

### 2.2 Animal Preparation for In Vivo Imaging

NDNF-Cre, CaMKIIα-Cre and PV-Cre mice are used in this study. All mice receive a total injection volume of 70 nl. NDNF-Cre animals are injected with pGP-AAV-syn-FLEX-Volton2-ST-WPRE virus into the visual cortex (from Bregma: AP -3.71 mm and -3.5 mm, ML +2.5 mm and 2.0 mm, DV −0.15 mm) [2]. CaMKIIα-Cre and PV-Cre mice are injected with pGP-AAV-syn-FLEX-Volton2-ST-WPRE virus in M1 (from Bregma: AP -1 mm, ML +1 mm, DV −0.15 mm) [2]. For pain management, sub-cutaneous injections of dexamethasone (.25 mg/kg) and buprenorphine (0.1 mg/kg) in saline are administered. After one month, a retro-orbital injection of Janelia Fluor 552 (JF552) is delivered and 4-24 hours later the mice are head-fixed beneath the objective. All surgeries and experiments performed in this work are approved by the University of Colorado Anschutz Medical Campus Institutional Animal Care and Use Committee (IACUC).

### 2.3 Data Processing

Data processing for the figures used the following steps. Structured illumination recordings are separated into 3 stacks (one for each phase of the grating pattern). The stacks are motion corrected individually using the VolPy implementation of NoRMCorre [19, 38]. For the mean OS-SIM reconstruction, as shown in Fig. 2B (ii), the motion corrected stacks for each phase of the grating pattern are then averaged in time to reduce contamination from shot noise. The resulting 3 frames, one for each phase of the grating pattern, are used in the OS-SIM algorithm resulting in a single averaged OS-SIM image for the entire recording. A FFT of the mean OS-SIM reconstruction (example in Figure 2C, ii) is obtained in ImageJ. For processing time traces from the structured illumination recording (Figures 4-6), the motion corrected stacks (obtained before averaging) are re-interleaved and a pWF reconstruction is obtained by applying a 3-frame moving average to each pixel. Finally, the pHiLo reconstruction is generated using the pWF and motion corrected image stacks as uniform illumination (I_UI_) and structured illumination (I_SI_), respectively. The alpha parameter used in HiLo reconstruction is attained by iteratively matching the FFT of the mean OS-SIM image with the FFT of the HiLo reconstruction. The mean OS-SIM image is used instead of the noisy OS-SIM image (Figure 2C, i) to better match the alpha parameter. ΔF/F time courses and spike templates used for characterization of temporal resolution are extracted using VolPy [19]. Traditional widefield recordings shown in Fig. 6, are obtained with all DMD mirrors turned on.

**Fig. 2.**
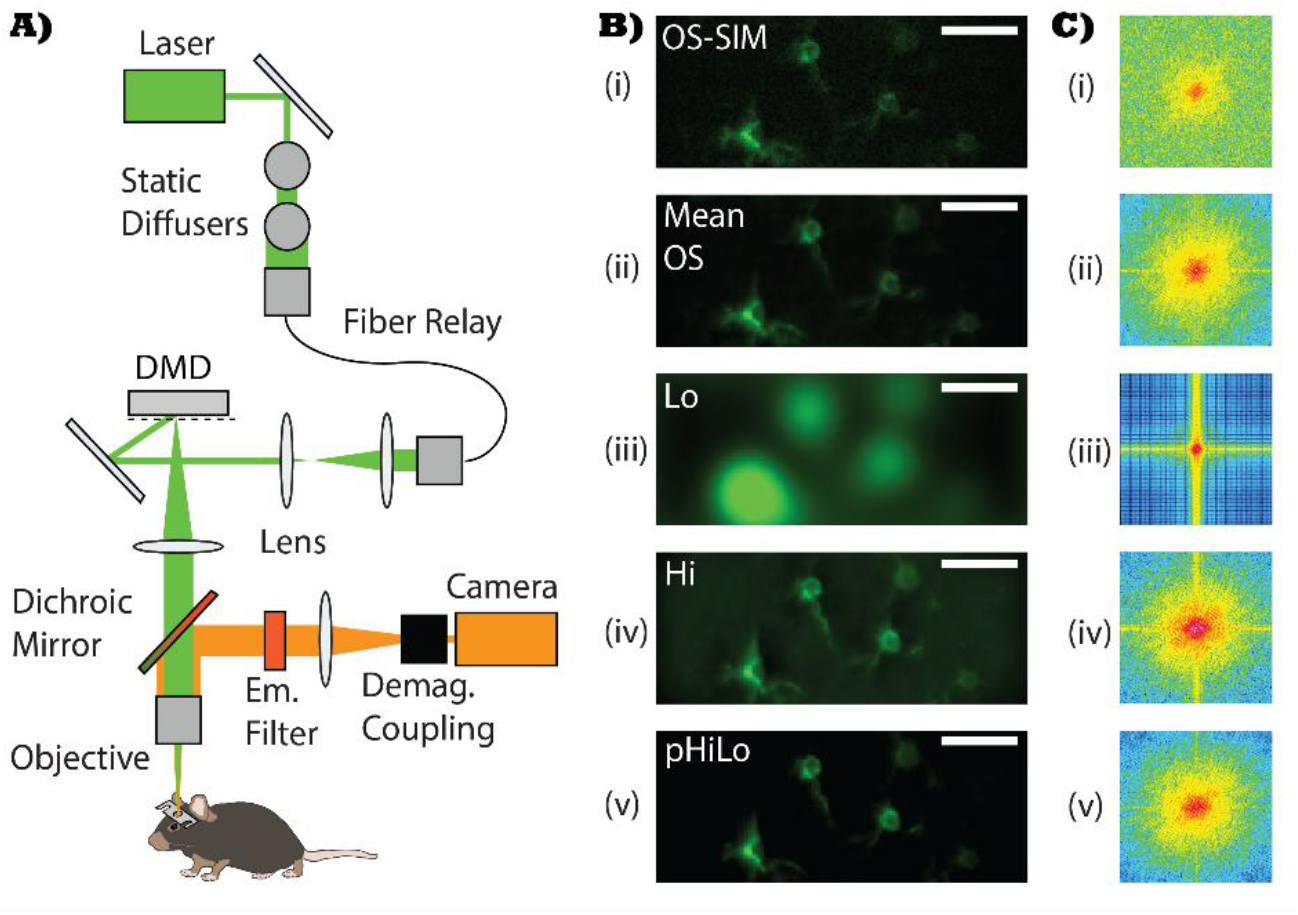
System Diagram and Image Reconstruction. **A**. Light emitted from a 532 nm laser passes through two diffusers and is fiber coupled to a multimode fiber to reduce coherent artifacts. The output of the fiber is collimated, patterned by a digital micromirror device and focused onto the sample using a 16x/0.8NA objective. Sample fluorescence is collected by the objective and reflected by a 567 nm shortpass dichroic mirror. Reflected fluorescence passes through a 568 nm long pass emission filter. A 0.35x demagnification coupling is used reduce the size of the image on the CMOS sensor. **B**. The noisy (i) and mean (ii) OS-SIM images. I_Lo_ (Lo) and I_Hi_ (Hi) images (iii-iv) used for pHiLo reconstruction (v). **C**. FFT images corresponding to frames in (B). The optimized pHiLo reconstruction is generated by adjusting α so that its FFT (v) matches that of the denoised OS-SIM reconstruction (ii).

## 3 Results

### 3.1 Characterization of Optical Sectioning Strength

Optical sectioning strength is characterized in data extracted from an NDNF-Cre mouse expressing Voltron2 in the visual cortex. To characterize the optical sectioning capability of the different techniques (Figure 3), we acquire widefield (Figure 3A), pHiLo (Figure 3B) and 2-photon (Figure 3C) recordings in the same field at different focal depths to obtain z-stacks. A 20 μm grating period was used for the pHiLo recording in Figure 3B. A region of information (ROI) is selected around a cell that is in-focus in the center frame of the z-stack. The total intensity within the ROI is measured at different depths and plotted in Figure 3D. For a 0.05 μm^-1^ spatial frequency illumination pattern, we calculate an optical sectioning strength of ∼14 μm using the Stokseth approximation [27]. The axial traces in Figure 3D are centered around the brightest plane for a cell that is in-focus at frame 0. The full width at half maximum (FWHM) for the pHiLo axial trace (∼ 30 μm) compared to the calculated sectioning strength (∼14 μm) is due to convolution with the size of the cell extending above/below this plane. For the 2P image, we calculate a sectioning strength of 1.26 μm as can be seen by the reduced FWHM of the plots shown in Figure 3D [39]. The decrease in FWHM for pHiLo compared to pWF, indicates axial sectioning capability. However, the increase in FWHM for pHiLo compared to 2-photon is due to the chosen grating frequency. To increase the optical sectioning capability for pHiLo, a higher-frequency grating pattern may be used. However, increasing grating frequency will inherently degrade the SNR of the sectioned reconstruction by reducing the modulation contrast of the grating pattern without increasing the overall signal collected.

**Fig. 3.**
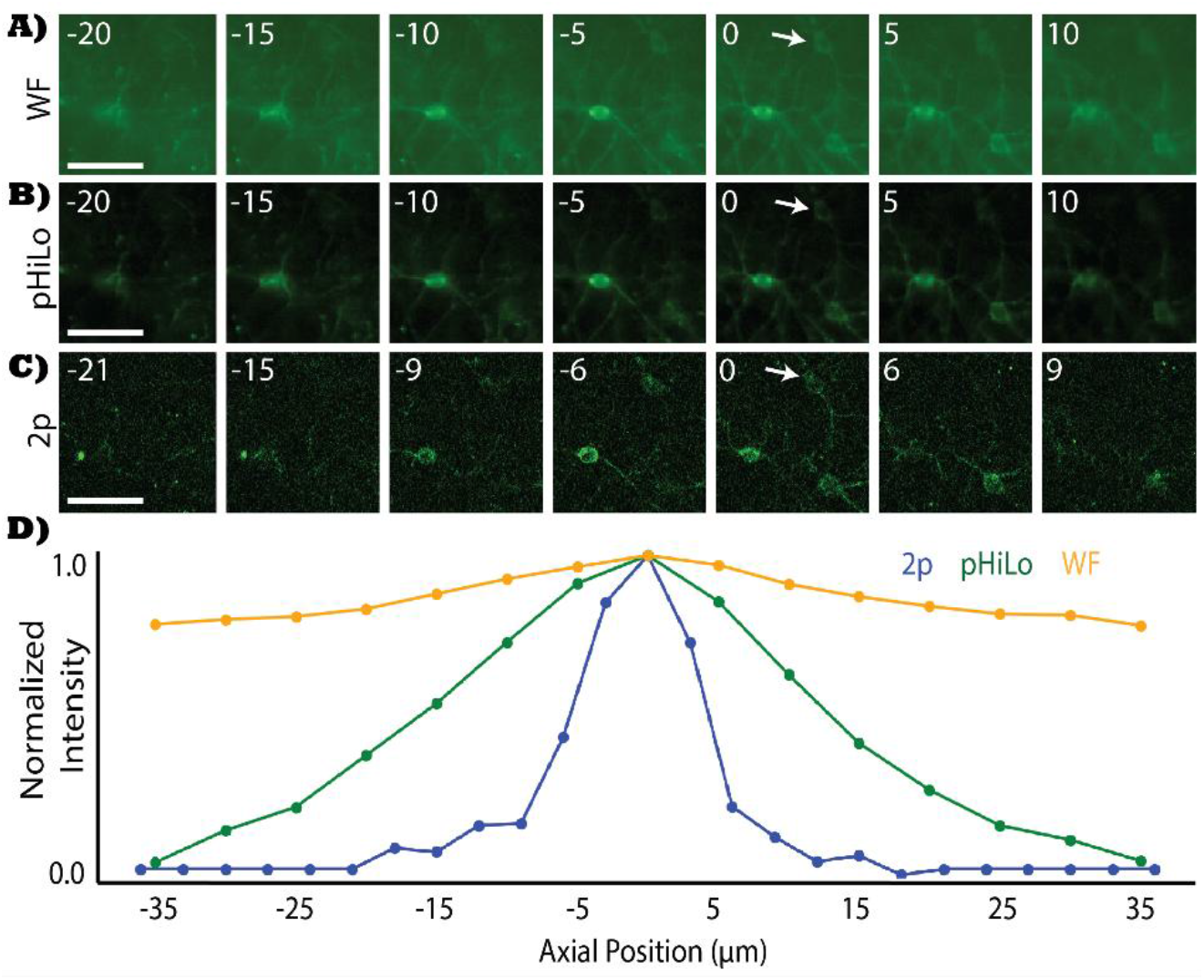
Comparison of widefield (WF), pseudo-HiLo (pHiLo) and 2-photon (2p) z-stacks extracted from the visual cortex of an awake mouse expressing Voltron2_552_. **A-C**. Frames were acquired with 5 μm steps (WF/pHiLo) and 3 μm steps (2p). A grating period of 20 μm was used for the pHiLo recordings. **D**. The total intensity for a region of interest (ROI) drawn around the cell (marked by the white arrow in A-C) was measured for each frame in the z-stack and plotted. The axial traces show optical sectioning of the pHiLo reconstruction in comparison with widefield and 2-photon.

The optical sectioning capability of the pHiLo reconstructions are illustrated in Figure 4 which compares intensity time courses from an in-focus cell and a cell that is defocused by 15 μm. Recordings for Figure 4 correspond to axial position -15 in Figure 3. Structured illumination images were acquired at 500 frames per second with a 20 μm grating period. Pseudo-WF (Figure 4A-B) and pHiLo (Figure 4B-C) reconstructions were obtained. Intensity time courses are plotted for the defocused cell (ROI 1) as well as a cell that is in-focus this plane (ROI 2). The optical sectioning of the pHiLo reconstruction removes the voltage signal of the out-of-focus cell (ROI 1) while preserving all the signal from the in-focus cell (ROI 2). It should also be noted that, while the data in Figure 4 was collected at 500 frames per second, clear transients are still visible despite the 3-frame moving average used in the pWF reconstruction.

**Fig. 4.**
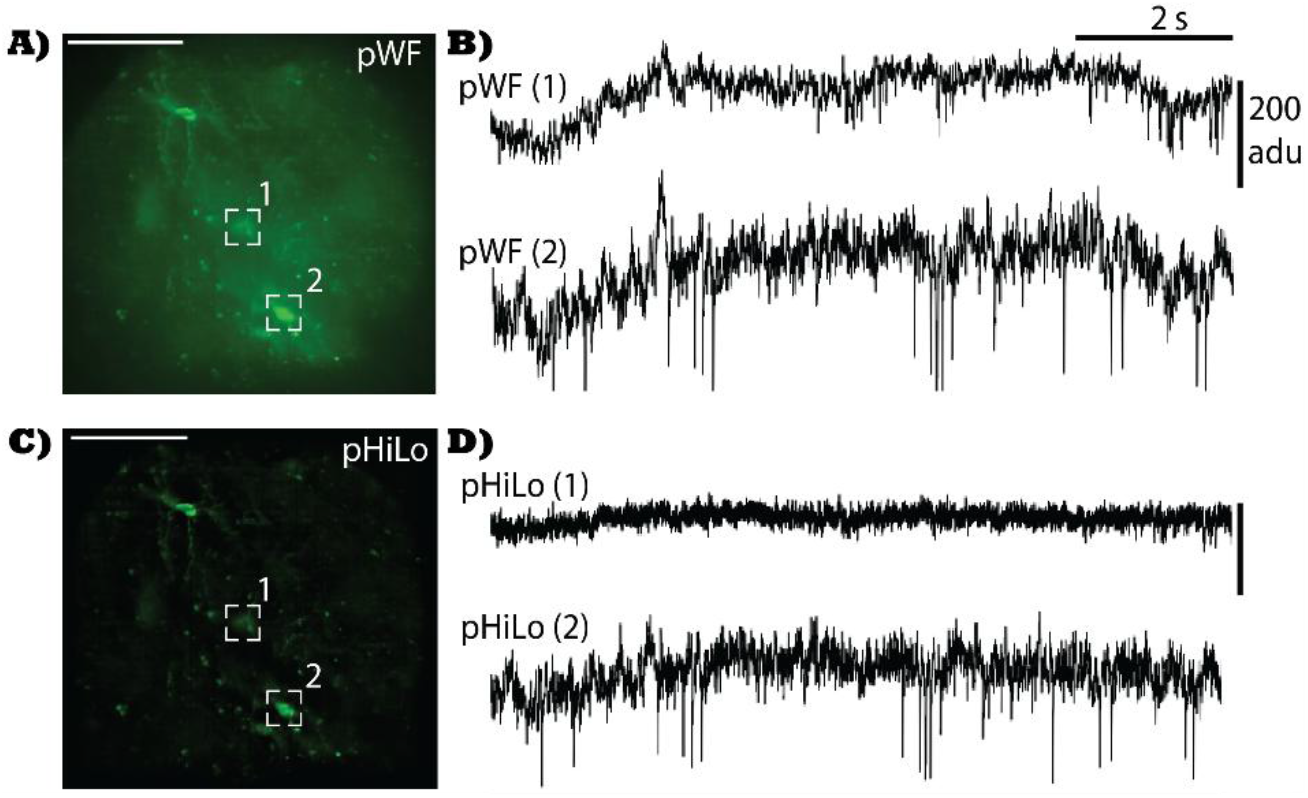
Voltage imaging with NDNF interneurons expressing Voltron2_552_ in the visual cortex using pseudo-HiLo (pHiLo). **A**. Average intensity projection (AIP) of the pseudo-Widefield (pWF) reconstruction. The scale bar is 100 μm. Data was collected with 100 mW/mm^2^ at 500 frames per second. **B**. Raw intensity time courses for the regions of information (ROI) shown in (A). ROI 1 corresponds to a cell that is defocused by 15 μm while ROI 2 is in focus at this plane. **C-D**. AIP and raw intensity time courses extracted from the pHiLo reconstruction. The ability for pHiLo to reduce background activity is indicated by the reduction in spiking magnitude for ROI 1 (the out-of-focus cell) that is not apparent for ROI 2 (the in-focus cell).

### 3.2 Background reduction in voltage recording using periodic structured illumination

To further evaluate the background reduction capabilities of this technique, data is extracted at 1200 frames per second from a CaMKIIα-Cre mouse expressing Voltron2 in the somatosensory cortex with a 25 μm grating pattern (Figure 5). Pseudo-Widefield and pHiLo reconstructions (Figure 5A) are performed and pixelwise correlation heatmaps (Figure 5B) are generated by calculating the Pearson correlation coefficient of the time course for each pixel with the mean time course for the full recording (sum of all pixels). ΔF/F time courses are then extracted from 5 ROIs to further evaluate background rejection capabilities.

**Fig. 5.**
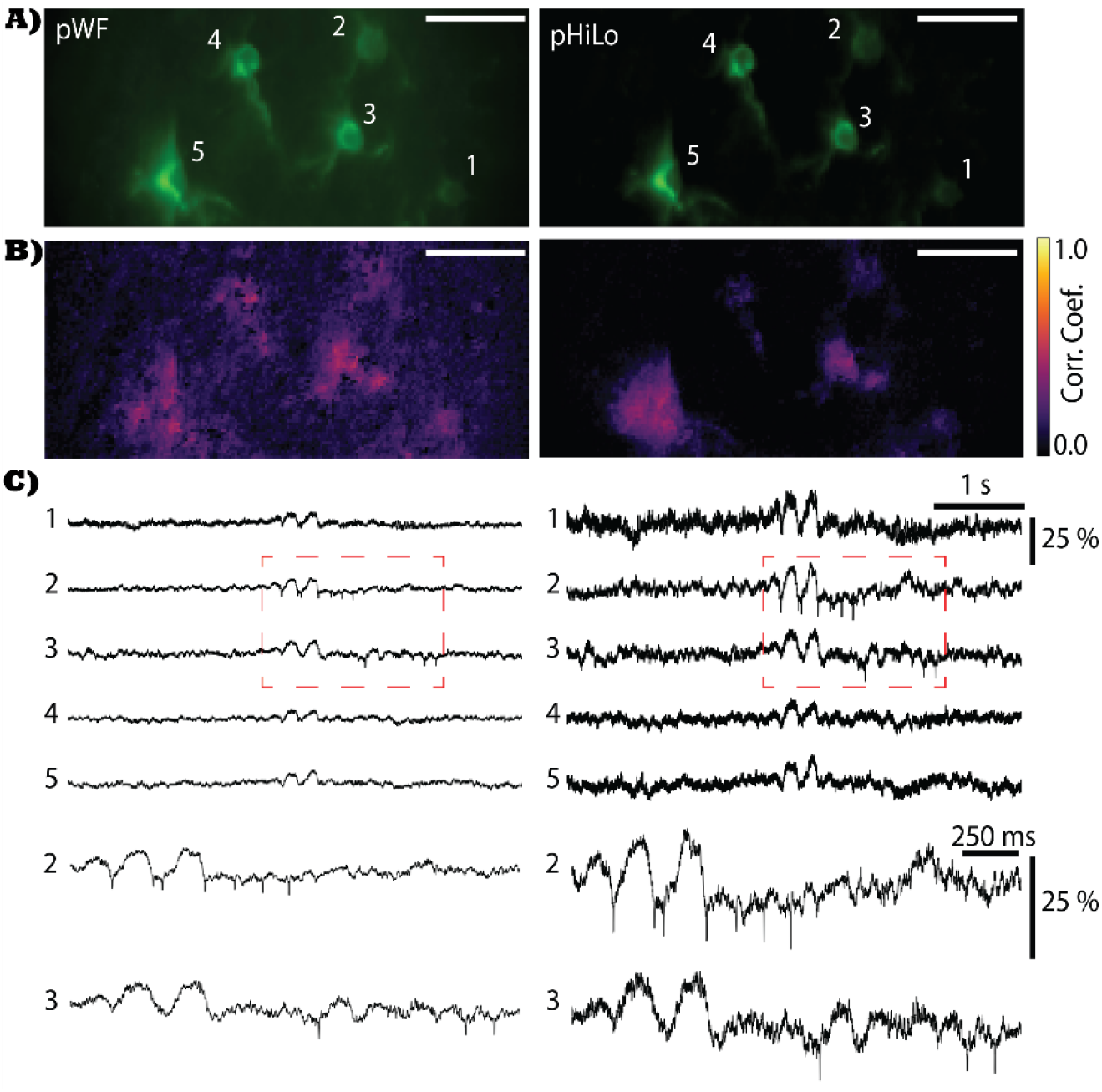
Voltage imaging in the somatosensory cortex using pseudo-HiLo (pHiLo). **A**. Average intensity projections of pseudo-Widefield (pWF, left) and pHiLo (right). The scale bar is 50 μm. Five cells expressing Voltron2_552_ in the somatosensory cortex are labelled in the pWF reconstruction. Data was recorded at 1200 frames per second using a grating period of 25 μm with a peak intensity of 100 mW/mm^2^ at the sample. **B**. Correlation heatmaps show the Pearson correlation coefficient (Corr. Coef.) between the intensity time course for each pixel with the mean intensity time course for all pixels. Out-of-focus fluorescence contamination in the pWF reconstruction (left) is indicated by the increased correlation between pixels that do not correspond to in-focus cells compared to the pHiLo reconstruction (right). **C**. Five second ΔF/F time courses extracted from the cells labelled in (A). A reduction in background contamination is indicated by the increase in ΔF/F for the pHiLo traces (right) compared to pWF (left). **D**. Two second activity windows for the red outlined portion of (C). Distinct voltage spikes can be seen in both reconstructions.

In Figure 5B, correlation heatmaps show a constriction in pixelwise correlation to in-focus cells for the pHiLo reconstruction that is not apparent in the pWF reconstruction. This visualization supports the ability for pHiLo to reduce fluctuations in out-of-focus transient activity compared to pWF. Additionally, the increase in ΔF/F percentage for pHiLo (Figure 5C-D) illustrates an increase in signal-to-background ratio compared to pWF. This increase is due to the reduction in resting baseline fluorescence (F) caused by the removal of out-of-focus background. A similar effect is apparent in Figure 6.

**Fig. 6.**
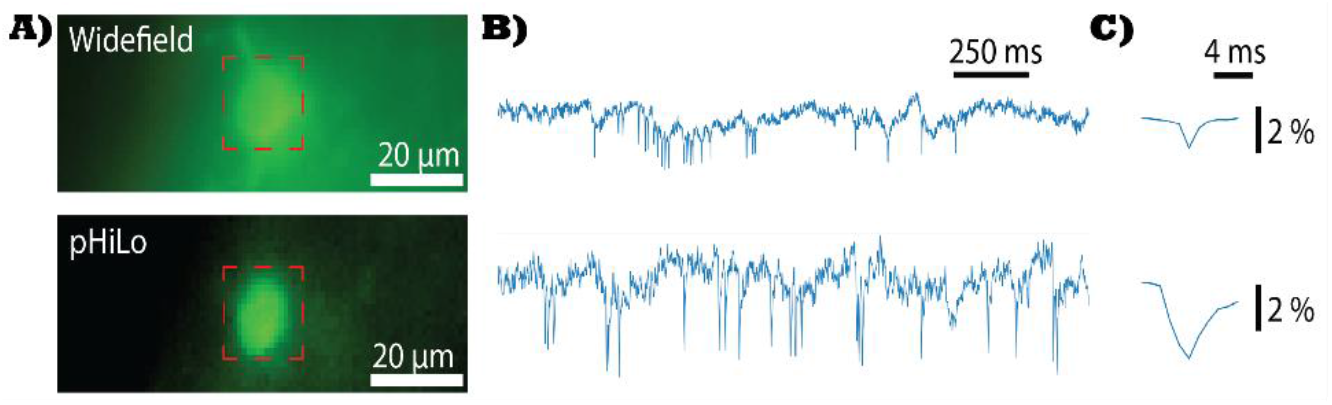
Comparison of widefield (top) and pHiLo (bottom) recordings taken from parvalbumin interneurons expressing Voltron2 in the somatosensory cortex. **A**. Average intensity projections of each recording. **B**. ΔF/F time courses extracted from the outlined ROIs in (A). **C**. VolPy extracted spike templates for each ROI time course. Data was recorded using 35 mW/mm^2^ intensity at 896 Hz (for the widefield recording) and 20 mW/mm^2^ intensity at 870 Hz with a grating period of 14 μm (for the pHiLo recording.

### 3.3 Characterization of Temporal Resolution

A PV-Cre mouse expressing Voltron2 in parvalbumin-positive interneurons in the somatosensory cortex was selected to compare the temporal resolution of pHiLo with traditional widefield recording (Figure 6). Data was first recorded using widefield illumination with 35 mW/mm^2^ intensity at 896 Hz and all DMD mirrors turned on (Figure 6, Top).

Structured illumination data was recorded with 20 mW/mm^2^ at 870 Hz, immediately after the widefield recording, using a grating period of 14 μm at the sample and pHiLo reconstruction was performed with an α of 1.0 (Figure 6, Bottom). ΔF/F time courses (Figure 6B) and spike templates (Figure 6C) are used to compare pHiLo with traditional widefield recording.

In Figure 6C, it is important to note the broadening of the pHiLo spike template compared to widefield recording with uniform illumination. Due to this broadening, the pHiLo technique requires a ∼3x higher sample rate than traditional widefield for cell types that have fast spike rates (>100 spikes/second). For this reason, we chose a sampling rate of approximately 900 Hz to record spike trains from fast spiking PV interneurons.

## 4. Discussion

The optical sectioning capabilities of pHiLo, compared to widefield, are indicated by the decrease in FWHM (Figure 3D), the reduction in magnitude for defocused cell activity (Figure 4), the constriction of pixelwise correlation to in-focus cell activity (Figure 5B) and the increase in ΔF/F magnitude for in-focus ROIs (Figure 5C-D, Figure 6B). While these metrics clearly convey the optical sectioning capabilities of pHiLo, the broadening of the pHiLo spike template in Figure 6C illustrates a tradeoff between signal-to-background ratio and temporal resolution. Despite this, the clear spikes in Figure 4B and 4D show that pHiLo can be effective at 500 Hz for detecting voltage spikes in NDNF interneurons. Our previous work indicates that interleaved OS-SIM techniques that rely on squared subtractions are not ideal for detecting GCaMP8f transients at 500 Hz due to distortions caused by the OS-SIM algorithm [1, 26]. By performing a HiLo reconstruction with each raw SI frame and the pWF (the average of the raw SI frame with the preceding and succeeding frame) at each time point, pHiLo overcomes this limitation and enables successful Voltron2 recording at 500-1200 frames per second. By demonstrating voltage imaging with pHiLo, we show that optically sectioned reconstructions of voltage activity may be obtained using only periodic structured illumination with simple stripe arrays. While additional techniques have been demonstrated for voltage imaging with patterned illumination [20, 35], it is our hope that the simplicity of pHiLo and the ease at which it may be combined with open-source data processing packages such as VolPy [19], enables its future use by neuroscience researchers whose studies require voltage imaging in awake/behaving animals.

## 5. Conclusion

In this study, we used periodic structured illumination to record voltage spike activity in awake, head-fixed mice expressing Voltron2 in various cell types and brain regions. We found that pHiLo could effectively remove background contamination from out-of-focus cells and allow the successful detection of voltage spikes in recordings taken from 500-1200 frames per second. By evaluating z-stacks extracted from NDNF-Cre neurons in the visual cortex, we found that pHiLo provides improved optical sectioning capabilities compared to widefield. Recording from PV interneurons in the somatosensory cortex, we show that pHiLo reconstructions cause a 3x broadening in extracted spike templates but provides an increase in spike ΔF/F.

The motivation for this work stems from the recent development of μLED stripe arrays designed to provide excitation with periodic structured illumination onboard miniature microscopes [16, 17]. In previous work, we demonstrated the capabilities of these stripe arrays as an OS-SIM light source [17]. Due to limitations of OS-SIM for functional imaging in awake mice, we developed pHiLo as an alternative reconstruction technique with increased resilience to distortions from motion in awake animals which can contaminate temporal processes [26]. In this work, we demonstrate that the optical sectioning benefits of pHiLo may be extended to voltage imaging experiments in awake mice.

## Acknowledgements

Portions of this work were presented at Optica Biophotonics Congress: Optics and the Brain, BM1B.3, “Voltage Imaging in Vivo Using Periodic Structured Illumination with Pseudo-HiLo Reconstruction”

## Data availability

All data and software presented in this paper may be obtained upon reasonable request.

## References

1. Y. Zhang, M. Rozsa, Y. Liang, et al., “Fast and sensitive GCaMP calcium indicators for imaging neural populations,” Nature 615, 884–891 (2023).

2. A. S. Abdelfattah, J. Zheng, A. Singh, et al., “Sensitivity optimization of a rhodopsin-based fluorescent voltage indicator,” Neuron 111, 1547–1563 e1549 (2023).

3. A. S. Abdelfattah, T. Kawashima, A. Singh, et al., “Bright and photostable chemigenetic indicators for extended in vivo voltage imaging,” Science 365, 699–704 (2019).

4. M. Ma, F. Simoes de Souza, G. L. Futia, et al., “Sequential activity of CA1 hippocampal cells constitutes a temporal memory map for associative learning in mice,” Curr Biol 34, 841–854 e844 (2024).

5. V. P. Sotskov, N. A. Pospelov, V. V. Plusnin, and K. V. Anokhin, “Calcium Imaging Reveals Fast Tuning Dynamics of Hippocampal Place Cells and CA1 Population Activity during Free Exploration Task in Mice,” Int J Mol Sci 23 (2022).

6. M. Kannan, G. Vasan, S. Haziza, et al., “Dual-polarity voltage imaging of the concurrent dynamics of multiple neuron types,” Science 378, eabm8797 (2022).

7. V. Villette, M. Chavarha, I. K. Dimov, et al., “Ultrafast Two-Photon Imaging of a High-Gain Voltage Indicator in Awake Behaving Mice,” Cell 179, 1590–1608 e1523 (2019).

8. F. P. Brooks, 3rd, D. Gong, H. C. Davis, et al., “Photophysics-informed two-photon voltage imaging using FRET-opsin voltage indicators,” Sci Adv 11, eadp5763 (2025).

9. J. Platisa, X. Ye, A. M. Ahrens, et al., “High-speed low-light in vivo two-photon voltage imaging of large neuronal populations,” Nat Methods 20, 1095–1103 (2023).

10. X. Lu, Y. Wang, Z. Liu, et al., “Widefield imaging of rapid pan-cortical voltage dynamics with an indicator evolved for one-photon microscopy,” Nat Commun 14, 6423 (2023).

11. J. Wu, Y. Liang, S. Chen, et al., “Kilohertz two-photon fluorescence microscopy imaging of neural activity in vivo,” Nat Methods 17, 287–290 (2020).

12. R. R. Sims, I. Bendifallah, C. Grimm, et al., “Scanless two-photon voltage imaging,” Nat Commun 15, 5095 (2024).

13. Z. Liu, X. Lu, V. Villette, et al., “Sustained deep-tissue voltage recording using a fast indicator evolved for two-photon microscopy,” Cell 185, 3408–3425 e3429 (2022).

14. Q. T. K. Lai, G. G. K. Yip, J. Wu, et al., “High-speed laser-scanning biological microscopy using FACED,” Nat Protoc 16, 4227–4264 (2021).

15. F. Phil Brooks, 3rd, H. C. Davis, J. D. Wong-Campos, and A. E. Cohen, “Optical constraints on two-photon voltage imaging,” Neurophotonics 11, 035007 (2024).

16. O. D. Supekar, A. Sias, S. R. Hansen, et al., “Miniature structured illumination microscope for in vivo 3D imaging of brain structures with optical sectioning,” Biomed Opt Express 13, 2530–2541 (2022).

17. V. Kumar, K. Behrman, F. Speed, et al., “MicroLED light source for optical sectioning structured illumination microscopy,” Opt Express 31, 16709–16718 (2023).

18. P. Zhou, S. L. Resendez, J. Rodriguez-Romaguera, et al., “Efficient and accurate extraction of in vivo calcium signals from microendoscopic video data,” Elife 7 (2018).

19. C. Cai, J. Friedrich, A. Singh, et al., “VolPy: Automated and scalable analysis pipelines for voltage imaging datasets,” PLoS Comput Biol 17, e1008806 (2021).

20. S. Xiao, W. J. Cunningham, K. Kondabolu, et al., “Large-scale deep tissue voltage imaging with targeted-illumination confocal microscopy,” Nat Methods 21, 1094–1102 (2024).

21. M. A. Lauterbach, E. Ronzitti, J. R. Sternberg, et al., “Fast Calcium Imaging with Optical Sectioning via HiLo Microscopy,” PLoS One 10, e0143681 (2015).

22. D. Lim, T. N. Ford, K. K. Chu, and J. Mertz, “Optically sectioned in vivo imaging with speckle illumination HiLo microscopy,” J Biomed Opt 16, 016014 (2011).

23. R. Cao, Y. Li, Y. Zhou, et al., “Dark-based optical sectioning assists background removal in fluorescence microscopy,” Nat Methods (2025).

24. M. Hupfel, A. Yu Kobitski, W. Zhang, and G. U. Nienhaus, “Wavelet-based background and noise subtraction for fluorescence microscopy images,” Biomed Opt Express 12, 969–980 (2021).

25. Y. Zhang, G. Zhang, X. Han, et al., “Rapid detection of neurons in widefield calcium imaging datasets after training with synthetic data,” Nat Methods 20, 747–754 (2023).

26. F. Speed, C. A. Saladrigas, A. Teel, et al., “High-speed in vivo calcium recording using structured illumination with self-supervised denoising,” Opt. Continuum 3, 2044–2059 (2024).

27. M. A. Neil, R. Juskaitis, and T. Wilson, “Method of obtaining optical sectioning by using structured light in a conventional microscope,” Opt Lett 22, 1905–1907 (1997).

28. J. Mertz, “Optical microscopy with planar or structured illumination,” Nat. Methods 8, 811–819 (2011).

29. P. T. Brown, R. Kruithoff, G. J. Seedorf, and D. P. Shepherd, “Multicolor structured illumination microscopy and quantitative control of polychromatic light with a digital micromirror device,” Biomed Opt Express 12, 3700–3716 (2021).

30. H. Zhang, Y. Zhu, L. Jin, et al., “Recent Advances in Structured Illumination Microscopy: From Fundamental Principles to AI-Enhanced Imaging,” Small Methods 9, e2401616 (2025).

31. Z. Li, Q. Zhang, S. W. Chou, et al., “Fast widefield imaging of neuronal structure and function with optical sectioning in vivo,” Sci Adv 6, eaaz3870 (2020).

32. R. Shi, Y. Li, and L. Kong, “High-speed volumetric imaging in vivo based on structured illumination microscopy with interleaved reconstruction,” J Biophotonics 14, e202000513 (2021).

33. T. Knopfel and C. Song, “Optical voltage imaging in neurons: moving from technology development to practical tool,” Nat Rev Neurosci 20, 719–727 (2019).

34. T. N. Ford, D. Lim, and J. Mertz, “Fast optically sectioned fluorescence HiLo endomicroscopy,” Biomed Opt 17, 021105 (2012).

35. V. Parot, C. Sing Long, Y. Adam, et al., “Compressed Hadamard microscopy for high-speed optically sectioned neuronal activity recordings,” Journal of Physics D: Applied Physics 52, 144001 (2019).

36. S. Xiao, E. Lowet, H. J. Gritton, et al., “Large-scale voltage imaging in behaving mice using targeted illumination,” iScience 24, 103263 (2021).

37. C. A. Saladrigas, E. J. Miscles, V. Kumar, et al., “Wobulation in structured illumination microscopy using a tunable electrowetting prism,” in CLEO 2024, Technical Digest Series (Optica Publishing Group, 2024), SF3B.6.

38. E. A. Pnevmatikakis and A. Giovannucci, “NoRMCorre: An online algorithm for piecewise rigid motion correction of calcium imaging data,” J Neurosci Methods 291, 83–94 (2017).

39. W. R. Zipfel, R. M. Williams, and W. W. Webb, “Nonlinear magic: multiphoton microscopy in the biosciences,” Nat Biotechnol 21, 1369–1377 (2003).

